# Actomyosin regulation by Eph receptor signaling couples boundary cell formation to border sharpness

**DOI:** 10.1101/683631

**Authors:** Jordi Cayuso, Qiling Xu, Megan Addison, David G. Wilkinson

## Abstract

The segregation of cells with distinct regional identity underlies formation of a sharp border, which in some tissues serves to organise a boundary signaling centre. It is unclear whether or how border sharpness is coordinated with induction of boundary-specific gene expression. We show that forward signaling of EphA4 is required for border sharpening and induction of boundary cells in the zebrafish hindbrain, which we find both require kinase-dependent signaling, with a lesser input of PDZ domain-dependent signaling. We find that boundary-specific gene expression is regulated by myosin II phosphorylation, which increases actomyosin contraction downstream of EphA4 signaling. Myosin phosphorylation leads to nuclear translocation of Taz, which together with Tead1a is required for boundary marker expression. Since actomyosin contraction maintains sharp borders, there is direct coupling of border sharpness to boundary cell induction that ensures correct organisation of signaling centres.

## Introduction

During embryo development, sharp borders form at the interface of adjacent tissues and between domains within tissues that have a different regional identity. These borders are generated by cell segregation mechanisms that establish and maintain a precise organisation of tissues (Batlle and Wilkinson, 2012; Dahmann et al., 2011; Fagotto, 2014). At some borders, a distinct boundary cell population is induced which serves as a signaling centre that regulates the patterning of cell differentiation within the tissue. The formation of a sharp and straight border enables such boundary signaling cells to be correctly organised (Dahmann and Basler, 1999). It remains unclear whether or how the induction of a signaling centre is coordinated with border sharpening. In principle, border sharpening and formation of boundary signaling cells may involve parallel mechanisms that are not directly linked. However, studies of the vertebrate hindbrain found that Eph receptor and ephrin signaling is required both for border sharpening and the formation of boundary cells (Cooke et al., 2005; Terriente et al., 2012; Xu et al., 1995), raising the possibility that there is a mechanistic link.

Eph receptor and ephrin signaling has major role in cell segregation and border sharpening in many tissues in vertebrates (Batlle and Wilkinson, 2012; Cayuso et al., 2015; Fagotto et al., 2014). Eph receptors comprise a large family of receptor tyrosine kinases that are activated upon binding to their membrane-bound ephrin ligands. Members of the EphA subclass bind to the GPI-anchored ephrinA ligands, whereas EphB receptors bind to transmembrane ephrinB ligands; an exception is EphA4 which binds to ephrinA and specific ephrinB family members (Gale et al., 1996). Upon interacting through cell-cell contact, Eph receptor and ephrin proteins are clustered and this activates signal transduction through both components, termed forward and reverse signaling, respectively (Klein, 2012; Pasquale, 2008). For Eph receptors, this involves kinase-dependent signaling that activates multiple intracellular pathways. In addition, signaling is mediated by a motif at the C-terminus of Eph receptors that binds to PDZ domain proteins. In the case of ephrinB proteins, signaling occurs through phosphorylation of conserved tyrosine residues by cytoplasmic kinases, and also through interaction of PDZ domain proteins.

Eph receptors and ephrins that have a high affinity for each other are expressed in complementary domains in many tissues (Cayuso et al., 2015; Gale et al., 1996; Rohani et al., 2014), such that activation of forward and reverse signaling occurs at the interface. Eph receptor and ephrin signaling can drive cell segregation and border sharpening through multiple mechanisms that likely depend upon whether the tissue is epithelial or mesenchymal: by decreasing cell-cell adhesion (Fagotto et al., 2013; Solanas et al., 2011), by increasing cortical tension (Calzolari et al., 2014; Canty et al., 2017), or by triggering cell repulsion (Poliakov et al., 2008; Rohani et al., 2011; Taylor et al., 2017; Wu et al., 2019). In addition, Eph-ephrin signaling has been found to regulate cell differentiation in a number of tissues (reviewed by (Laussu et al., 2014; Wilkinson, 2014)). The regulation of cell differentiation and segregation may be distinct and context-dependent functions. Alternatively, Eph-ephrin signaling could couple cell specification to maintenance of their organisation. For most tissues, it is unclear whether such coupling occurs, but a potential example is the formation of boundaries in the vertebrate hindbrain.

The hindbrain is subdivided into segments, termed rhombomeres (r1-r7), each with a distinct anteroposterior identity and demarcated by borders across which cell intermingling is restricted (Fraser et al., 1990). These borders are initially fuzzy and then sharpened through the regulation of cell identity (Addison et al., 2018; Wang et al., 2017) in combination with cell segregation driven by Eph-ephrin signaling (Cooke et al., 2005; Kemp et al., 2009; Xu et al., 1995; Xu et al., 1999). Boundary cells are induced to form at segment borders (Guthrie and Lumsden, 1991) and express specific molecular markers that distingish them from non-boundary cells (Cheng et al., 2004; Cooke et al., 2005; Heyman et al., 1995; Xu et al., 1995). In zebrafish, these include the Notch modulator, *rfng*, which by inhibiting neurogenesis promotes the maintenance of boundary cells (Cheng et al., 2004). Boundary cells have been shown to act as a signaling centre that organises spatially-restricted neurogenesis within hindbrain segments in zebrafish (Gonzalez-Quevedo et al., 2010; Terriente et al., 2012). Several Eph receptors are segmentally expressed in the hindbrain, in a complementary pattern to ephrinBs that they have high affinity for: *ephA4* in r3 and r5 is complementary to *ephrinB3* in r2, r4 and r6; *ephB4* in r5 and r6 is complementary to *ephrinB2* in r1, r4 and r7. Disruption of Eph receptor or ephrin function leads to a decrease both in the sharpening of segment borders and in the expression of boundary markers (Cooke et al., 2005; Terriente et al., 2012; Xu et al., 1995). These findings raise the questions of how Eph-ephrin signaling leads to boundary cell formation and whether this involves distinct pathways from border sharpening.

We set out to dissect mechanisms of signaling that underlie border sharpening and boundary cell specification in the zebrafish hindbrain. EphA4 and ephrinB3 act as a signaling pair since knockdown of either component disrupts the same segment boundaries (Terriente et al., 2012). We find that boundary cell markers are expressed in *epha4*-expressing cells and are up-regulated by forward signaling. By creating a series of truncation and point mutants in *epha4*, we show that kinase-dependent and PDZ domain-dependent signaling both contribute to regulation of border sharpening and boundary-specific gene expression. We find that boundary marker expression is regulated by myosin II phosphorylation that occurs downstream of EphA4 activation and increases mechanical tension at segment borders. Mechanotransduction that induces boundary marker expression is mediated by nuclear translocation of Taz. The regulation of actomysosin contraction by Eph signaling thus couples the maintenance of sharp borders and induction of a boundary signaling centre.

## Results

### Boundary marker expression occurs in ephA4-expressing cells

Since *epha4* (*epha4a*) is expressed in r3 and r5 (Xu et al., 1995), and *ephrinb3* (*efnb3b*) in r2, r4 and r6 (Chan et al., 2001), this Eph-ephrin pair interacts at all borders of r3 and r5. Due to the bidirectionality of activation, knockdown of either component will lead to loss of both Eph and ephrin activation, and it is therefore not possible to deduce whether forward and/or reverse signaling regulates boundary marker expression. A clue can come from determining whether boundary cells form in *epha4*-expressing cells, *ephrinb3*-expressing cells, or both. To address this, we carried out *in situ* analysis using the hybridisation chain reaction (HCR) which enables sensitive fluorescent detection of multiple transcripts (Choi et al., 2016). We found that *rfng* expression which marks hindbrain boundary cells occurs in *epha4*-expressing cells at the borders of r3 and r5 (Fig.1A-C). *rfng* expression is also detected in a few cells that are not expressing *epha4*, which are most consistently found at the lateral edge of the r5/r6 border (arrow in Fig.1B, C). We also analysed expression of *wnt1*, which is expressed in the roof plate and in the dorsal part of hindbrain boundaries (Fig.1D). We found that *wnt1* expression in boundaries also occurs predominantly in *epha4*-expressing cells (Fig.1E, F). Boundary cell formation thus occurs in cells in which forward signaling is occurring. However, *rfng* expression also occurs in some cells that are not expressing *epha4*, which could reflect a role of reverse signaling, or a dynamic relationship between *epha4* and *rfng* gene expression.

**Figure 1:**
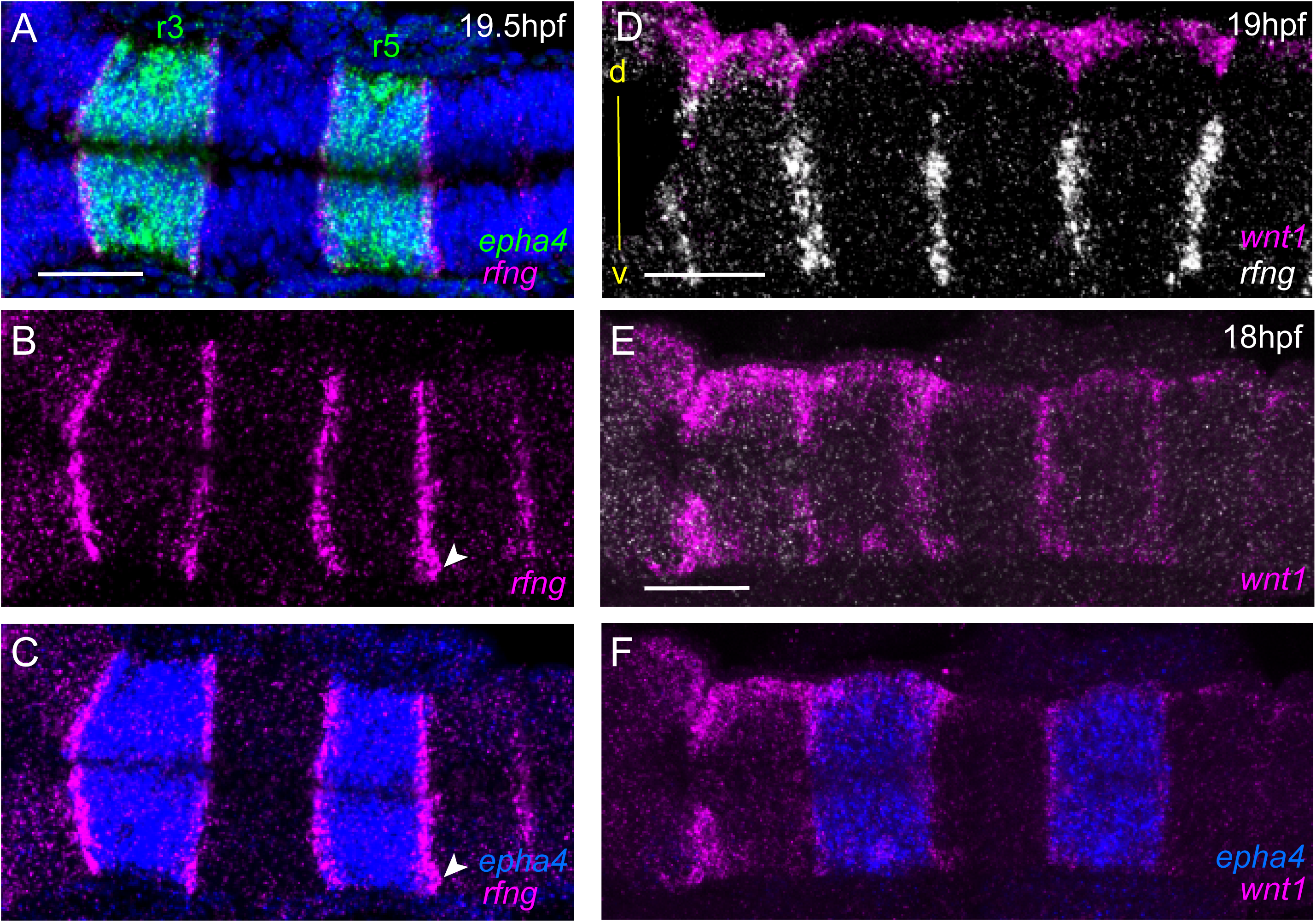
Boundary markers are expressed in epha4-expressing rhombomeres. **(A-C)** HCR stainings for *rfng* and *epha4*. *rfng* is expressed in *epha4*-expressing cells in rhombomeres r3 and r5, with the exception of a few *rfng-*expressing cells in r6 (arrowheads). **(D-F)** Boundary expression of *wnt1* is dorsal to *rfng* (D) and is coexpressed with epha4 (E, F). (A-C, E, F) are dorsal views, (F) is a lateral view, dorsal to the top. Anterior to the left in all panels. Scale bar: 50 μm.

### Border sharpening and boundary marker expression require forward signaling

Knockdown of *epha4* or *ephrinb3* leads to loss or decrease in expression of boundary cell markers at three borders where they interact (Fig.2A): r2/r3, r3/r4 and r5/r6 (Terriente et al., 2012); there is potential functional redundancy with *ephb4* and *ephrinb2* at the r4/r5 border (Chan et al., 2001; Cooke et al., 2001; Cooke et al., 2005). To test roles of different aspects of EphA4 signaling, we used CRISPR/Cas9 genome modification to create a series of zebrafish lines with point or truncation mutations, depicted in Fig.2B. The null mutant has a 4 bp deletion which terminates EphA4 protein within the ligand-binding domain. The truncation mutant terminates the protein at residue 651 (*epha4*^*Δ651*^), deleting most of the tyrosine kinase domain and all C-terminal domains, and thus completely lacks forward signaling but potentially still activates reverse signaling. The kinase-dead mutant (*ephA4*^*KD*^) replaces a lysine residue essential for kinase function with methionine. The *epha4*^*ΔPDZBD*^ mutant is truncated at residue 994 which removes the five C-terminal amino acids containing the PDZ binding domain (PDZBD) motif.

**Figure 2:**
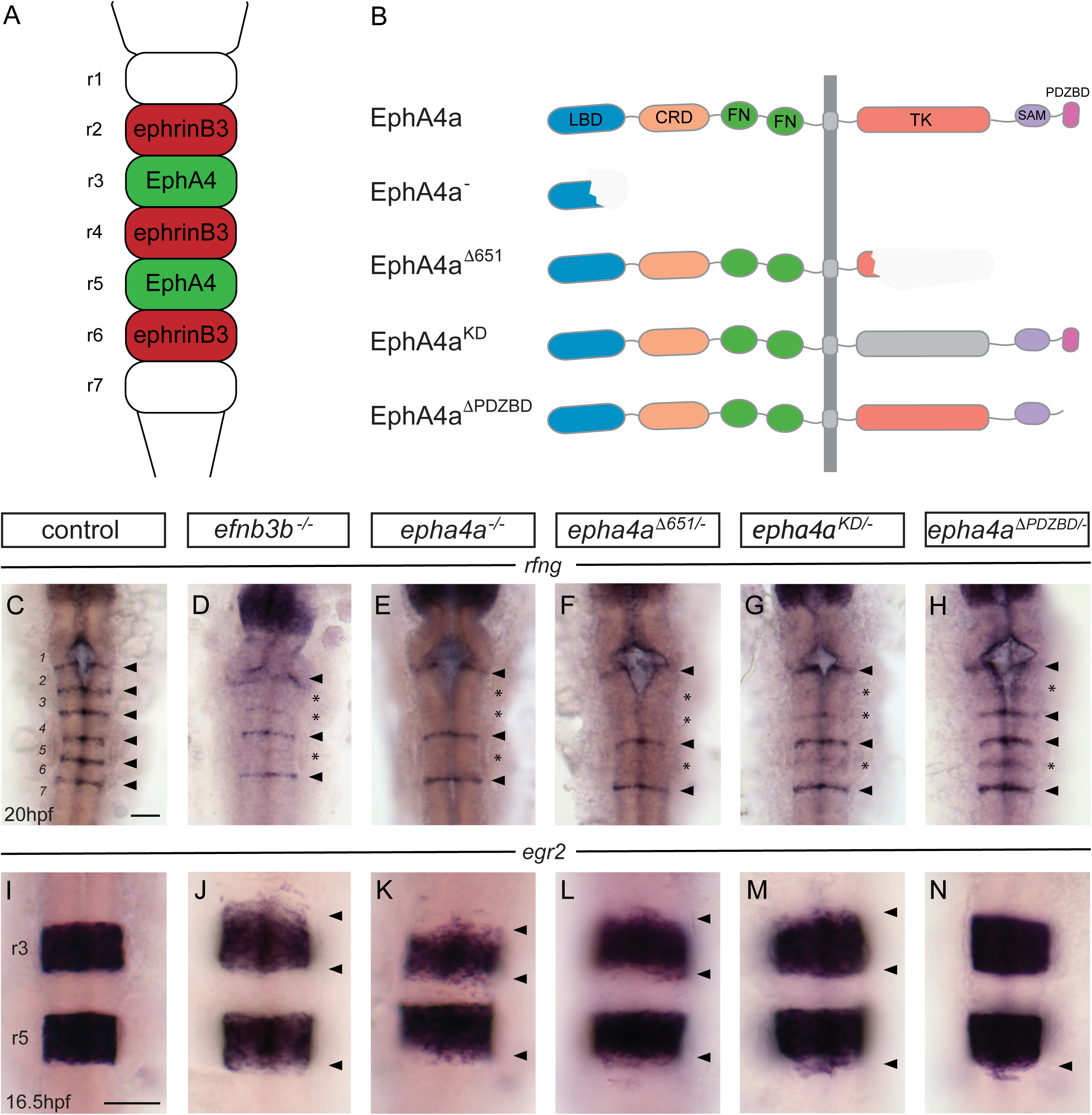
EphA4 forward signalling regulates boundary marker expression and cell segregation. **(A)** Schematic representation of the segmented expression of EphA4 and ephrinB3 in the hindbrain. **(B)** Schematic representation of the different mutant alleles of *ephA4a* generated for this study. The null allele contains an early truncation in the ligand binding domain. The *epha4*^*Δ651*^ allele lacks most of the cytosolic domain. The *ephA4*^*KD*^ allele contains a point mutation of a critical lysine in the tyrosine kinase domain. The *ephA4*^*ΔPDZBD*^ mutation consists of a C-terminal truncation that deletes the PDZ-binding domain. LBD – ligand binding domain; CRD – cysteine rich domain; FN – fibronectin repeat; TK – tyrosine kinase domain; SAM – sterile alpha motif; PDZBD – PDZ binding domain. **(C-H)** *rfng* is expressed at boundaries in control embryos (arrowheads) (C), but is reduced or absent (star) at specific boundaries in *ephrinB3*^*-/-*^ (D), *epha4*^*-/-*^ (E), *epha4*^*Δ651*^ (F), *epha4*^*KD*^ (G) and *epha4*^*ΔPDZBD*^ (H) mutants. **(I-N)** *egr2* expression in r3 and r5 has sharp borders in control embryos (I); border sharpening defects (arrowheads) are observed in *ephrinb3*^*-/-*^ (J), *epha4*^*-/-*^ (K), *epha4*^*Δ651*^ (L), *epha4*^*KD*^ (M) and *epha4*^*ΔPDZBD*^ (N) mutants. Dorsal views, anterior to the top in all panels. Scale bar: 50 μm.

We found that the null mutant of *epha4* has the same phenotype described previously for *epha4* knockdown, with loss of *rfng* expression at the r2/r3, r3/r4 and r5/r6 borders (Fig.2E; compare with wild type, Fig.2C). Furthermore, expression of other boundary markers, including *wnt1* and *sema3gb*, was disrupted at these borders (Suppl. Fig.1A-F). A milder disruption of *rfng* expression at the r2/r3, r3/r4 and r5/r6 borders is found in *ephrinB3* null mutant embryos (Fig.2D), likely reflecting some functional overlap with *ephrinB2* which is also a ligand for EphA4 (Cooke et al., 2005). The *epha4*^*Δ651*^ truncation mutant was found to have the same loss of *rfng* expression as *epha4* null mutants (Fig.2F), supporting the idea that boundary cell formation is dependent upon EphA4 forward signaling. However, since loss of the cytoplasmic domain of EphA4 might alter its activity as a ligand, this finding does not rule out a contribution of reverse signaling. To address whether kinase-dependent forward signaling is required, we analysed the *epha4*^*KD*^ mutant and found a major decrease, but not complete loss of *rfng* expression at the r2/r3, r3/r4 and r5/r6 borders (Fig.2G). The residual *rfng* expression at the r5/r6 border occurs in *epha4*-expressing cells (Suppl. Fig.1G, G’), arguing against the possibility that it is due to reverse signaling activated by *epha4*^*KD*^. The presence of some *rfng* expression at segment borders in the *epha4*^*KD*^ mutant suggests that kinase-independent signaling contributes to boundary cell formation. To test whether there is a parallel pathway involving signaling through PDZ domain proteins, we analysed the *epha4*^*ΔPDZBD*^ mutant. We found that there is a mild decrease in *rfng* expression at the r2/r3 and r5/r6 borders, though not at the r3/r4 border (Fig.2H). These findings are consistent with a contribution of PDZBD-dependent signaling to boundary cell formation.

Analysis of *egr2* expression in the e*pha4* null mutant revealed a decrease in sharpness of the r2/r3, r3/r4 and r5/r6 borders, with some *egr2*-expressing cells in the adjacent segments (Fig.2K; compare with control, Fig.2I). The sharpness of the same borders was disrupted in the *ephrinb3, epha4*^*Δ651*^ and *epha4*^*KD*^ mutants, though with fewer ectopic *egr2*-expressing cells compared with the null mutant (Fig.2J, L, M). In the *epha4*^*ΔPDZBD*^ mutant there was a decrease in sharpness at the r5/r6 border but only in 30% of the embryos at r2/r3 (Fig.2N). Taken together, these findings suggest that both kinase-dependent and PDZ domain-dependent pathways contribute to upregulation of boundary marker expression, with a stronger input of kinase signaling. There is a correlation between decreased border sharpness and decreased boundary marker expression, suggestive of a mechanistic link.

### Boundary marker expression is regulated by myosin phosphorylation

These findings raise the question of how EphA4 forward signaling leads to *rfng* expression at boundaries. EphA4 signaling regulates formation of an actin cable at boundaries, which is first detected at 15 hpf and has been implicated in maintenance of a straight border through actomyosin-dependent generation of cortical tension (Calzolari et al., 2014). Hindbrain boundary cells have a distinct shape from non-boundary cells, which is altered by knockdown of the myosin phosphatase regulator, *mypt1*, that leads to increased phosphorylation of myosin light chain (pMLC) and actomyosin contraction (Fig.3H) (Gutzman and Sive, 2010). Consistent with these findings, we found a higher level of pMLC co-localising with the actin cable at hindbrain borders (Fig.3A, B). Furthermore, pMLC was no longer detected at the r2/r3, r3/r4 and r5/r6 borders in *epha4* null mutants (Fig.3C, D). Surprisingly, we found that knockdown or transient CRISPR/Cas9-mediated knockout of *mypt1* leads to an increase in the level and width of *rfng* expression at hindbrain boundaries (Fig.3E-G). Likewise, *mypt1* knockdown leads to increased expression of *wnt1* and *sema3gb* at boundaries (Suppl. Fig.2A-D). We wondered whether the broader expression of *rfng* after *mypt1* knockdown occurs in regions of forward and/or reverse signaling. By carrying out *in situ* HCR we found that *rfng* expression spreads only into the *epha4*-expressing domain where forward signaling is occurring (Fig.3I-L). The increased boundary marker expression after *mypt1* knockdown suggests that actomyosin contraction regulates boundary cell formation. To test this, we treated embryos at different time intervals with blebbistatin, an inhibitor of myosin II ATPase activity. We found that blebbistatin treatment from 15 hpf onwards strongly disrupts the upregulation of *rfng* expression at boundaries, with a progressively milder effect on the expression when the treatment is started at later times, and no change detected when treated from 18 hpf (Fig.3M-P). Furthermore, disruption of actin polymerisation by treating embryos with latrunculinB leads to loss of *rfng* expression at boundaries (Suppl.Fig.2E, F).

**Figure 3:**
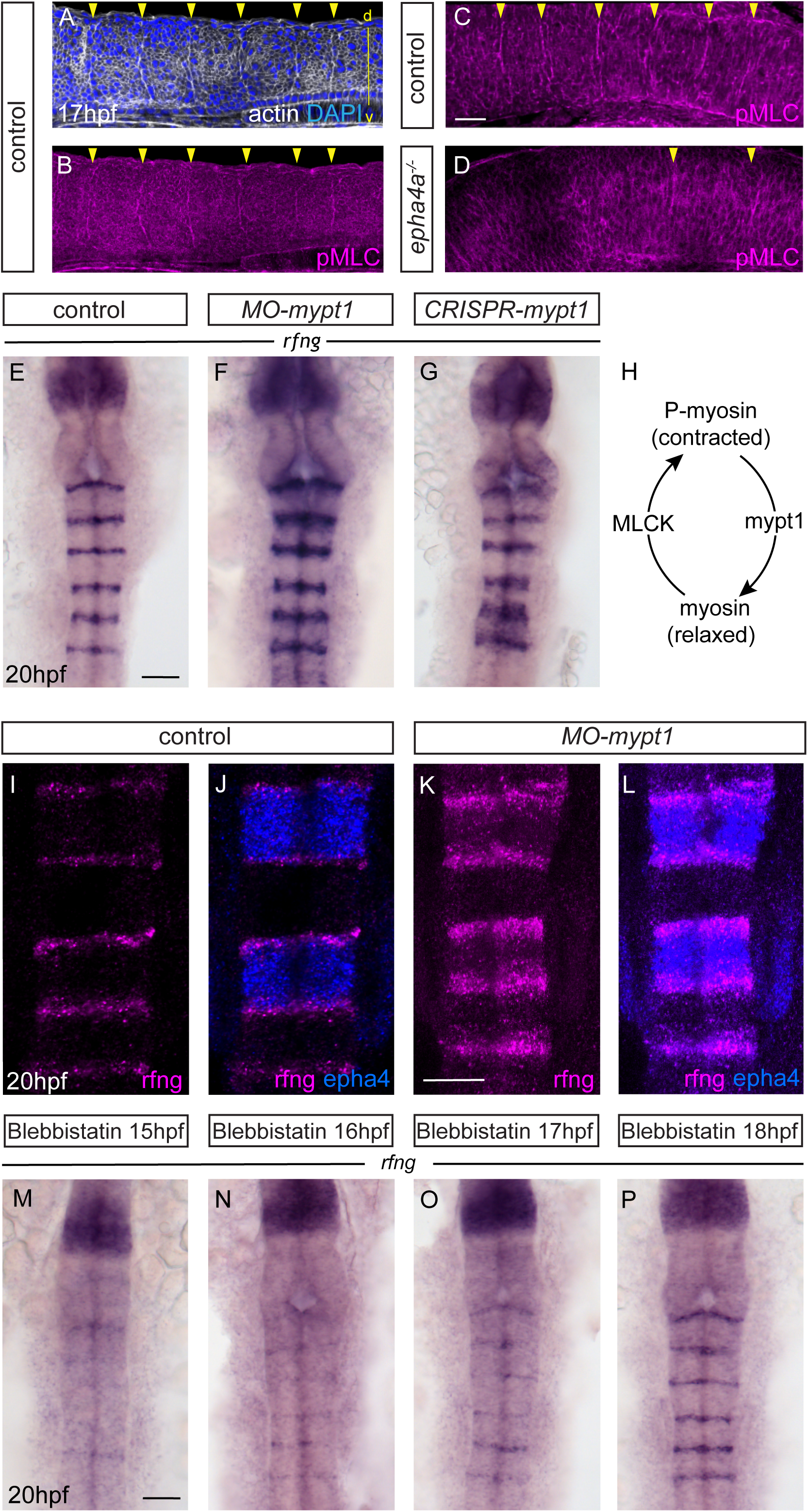
Actomyosin tension regulates boundary expression of *rfng*. **(A**, **B)** Immunostainings to detect actin (A) and pMLC (B) which co-localize at segment boundaries. **(C**, **D)** pMLC is detected at all boundaries in control embryos (C) and only at specific boundaries in *epha4*^*-/-*^ embryos (D). Lateral views, anterior to the left. **(E-G)** *rfng* expression is increased in *mypt1* knockdowns (F) and embryos injected with CRISPR/Cas9 against *mypt1* (G), compared to controls (E). **(H)** Depiction of mypt1 regulating actomyosin tension by dephosphorylating pMLC. **(I-L)** HCR stainings reveal that *rfng* is expressed in *epha4*-expressing cells in control embryos (I, J) and after knockdown of *mypt1* (K-L). **(M-P)** Myosin II inhibitor blebbistatin suppresses *rfng* transcription when treatment is initiated at 15 hpf (M), 16 hpf (N) or 17 hpf (O), but it is not affected when initiated at 18 hpf (P). (E-P) Dorsal views, anterior to the top. Scale bar: 50 μm.

These findings suggest that the induction of hindbrain boundary markers involves increased actomyosin contraction downstream of EphA4 forward signaling. We therefore wondered whether *mypt1* knockdown can rescue the decrease in *rfng* expression that occurs in *epha4* mutants. We found that *mypt1* knockdown in *epha4* null mutants rescues *rfng* expression at the r2/r3 and r3/4 borders, but not at the r5/r6 border (Fig.4C, H; wild type embryos in Fig.4A, F; quantitated in Fig.4K). This suggests that *mypt1* knockdown is increasing residual MLC phosphorylation at the r2/r3 and r3/4 borders in the *ephA4* null mutant, potentially due to other segmentally-expressed Eph receptors, whereas such compensation does not occur at the r5/r6 border. Intriguingly, *mypt1* knockdown rescues *rfng* expression at the r5/r6 border as well as the r2/r3 and r3/r4 borders in the *epha4*^*Δ651*^ and *epha4*^*KD*^ mutants (Fig.4D, E, I-K). This finding suggests that there is some compensation with mutants that have the EphA4 extracellular domain that does not occur when EphA4 protein is completely absent. *mypt1* knockdown rescues *rfng* expression at all hindbrain boundaries in *ephrinB3* null mutants (Fig.4B, G), consistent with residual EphA4 activation by other ephrins.

**Figure 4:**
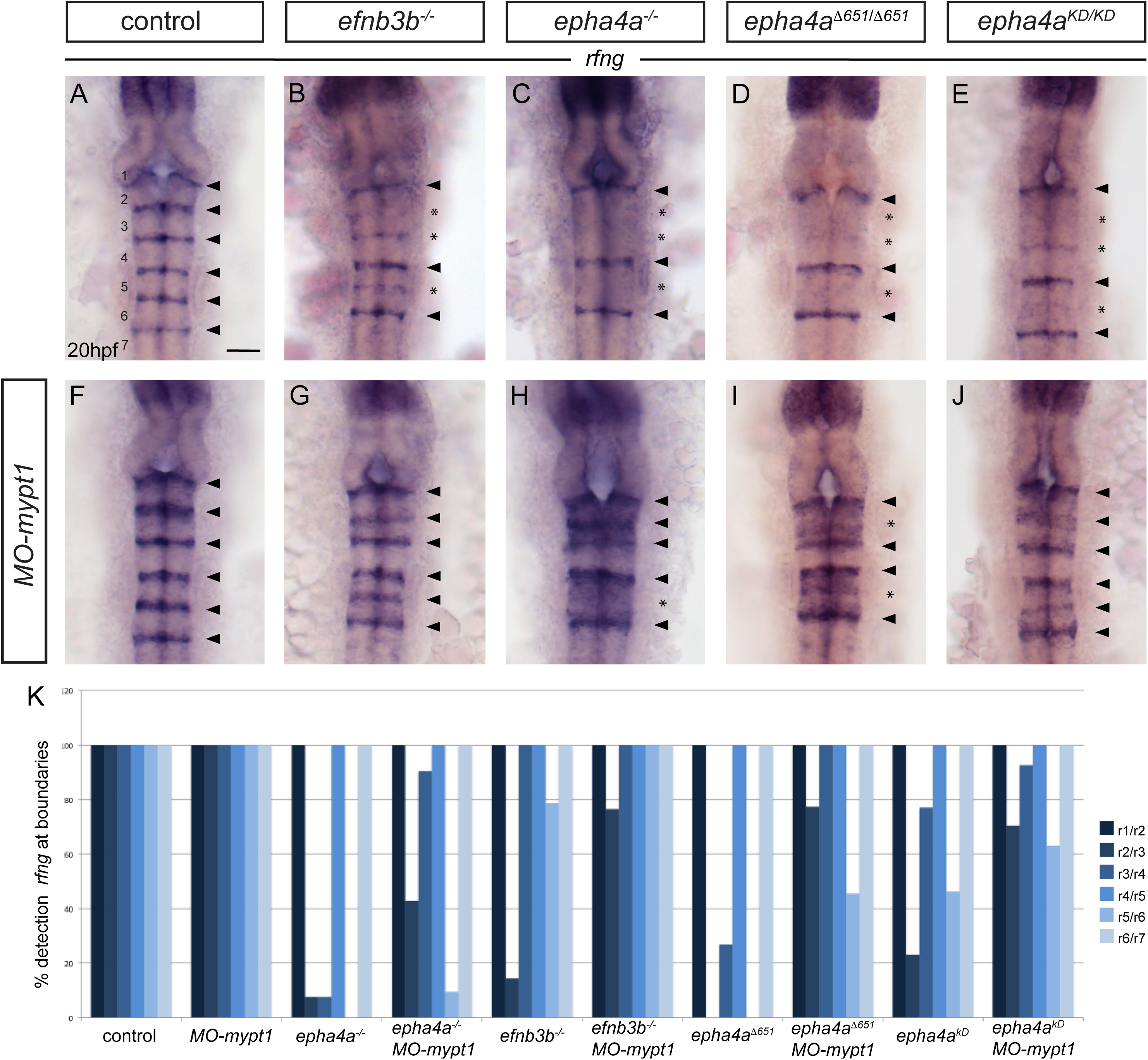
Increased tension selectively restores boundary expression of *rfng* in epha4 and ephrinb3 mutants. **(A-K)** Knockdown of *mypt1* increases *rfng* expression at hindbrain boundaries in control embryos (F) and restores *rfng* expression at specific boundaries in *ephrinb3*^*-/-*^ (G), *epha4*^*-/-*^ (H), *epha4*^*Δ651*^ (I) and *epha4*^*KD*^ (J) mutants compared to uninjected controls (A-E). (K) Percentage of embryos showing *rfng* expression at the different boundaries. Control (n=41); *MO-mypt1* (n=24); *epha4a*^*-/-*^ (n=13); *epha4a*^*-/-*^ *MO-mypt1* (n=42); *efnb3b*^-/-^ (n=14), *efnb3b*^-/-^ *MO-mypt1* (n=17), *epha4a*^*0651*^ (n=15), *epha4a*^*0651*^ *MO-mypt1* (n=22); *epha4a*^*KD*^ (n=13); *epha4a*^*KD*^ *MO-mypt1* (n=27). Dorsal views, anterior to the top. Arrowheads indicate normal boundary expression of *rfng*, while stars indicate reduction or absence of *rfng* expression at boundaries. Scale bar: 50 μm.

### taz and tead1a are required for boundary marker expression

Taken together, these findings suggest a model in which EphA4 forward signaling leads to actomyosin contraction that induces boundary marker expression. This raises the question of what pathway links mechanical tension to gene regulation at hindbrain boundaries. To address this, we carried out morpholino-mediated knockdowns of genes that have been implicated in mechanotransduction in other contexts. This screen revealed that knockdown of the *taz* gene disrupts boundary marker expression, including *rfng, wnt1* and *sema3gb* (Fig.5A, B, H; Suppl. A-D). To test the specificity of the gene knockdown, we carried out transient CRISPR/Cas9-mediated deletions of *taz* and found that this also leads to decreased *rfng* expression at boundaries (Fig.5C, H). In contrast, knockdown or knockout of the related *yap1* gene has no effect on boundary marker expression (Fig.5D, E, H). *taz* therefore has a non-redundant role in upregulation of boundary marker expression. The finding that *yap1* is not required may reflect relative expression levels, or differences in biochemical function of Taz and Yap (reviewed by (Callus et al., 2019)).

**Figure 5:**
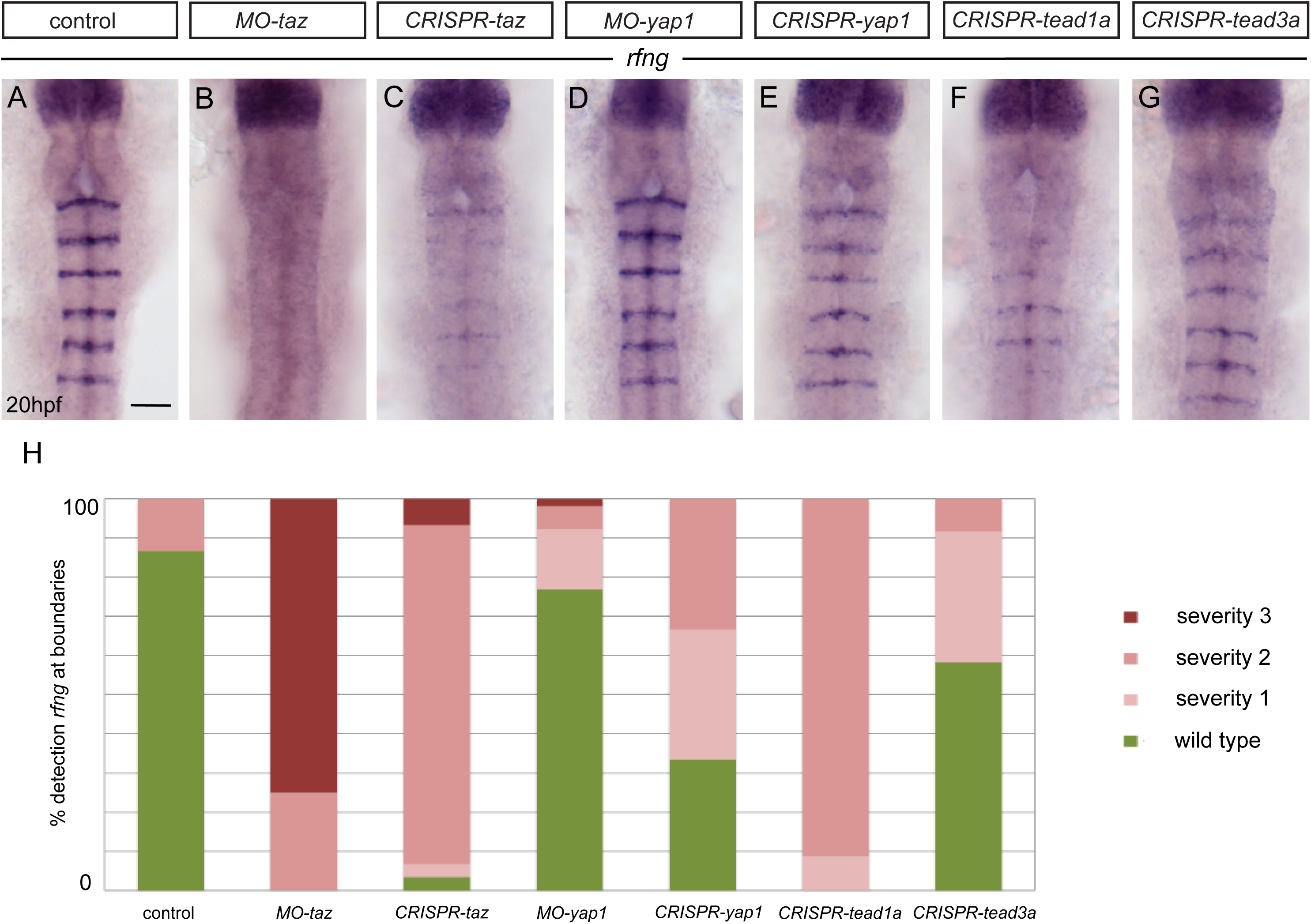
Taz and Tead1a are required for boundary expression of *rfng*. **(A-H)** Hindbrain boundary expression of *rfng* is reduced in *taz* knockdowns (B), and in *taz* (C) and *tead1a* (F) transient knockouts compared to controls (A), while *yap1* knockdown (D) and *yap1* (E) and *tead3a* (G) transient knockouts have normal *rfng* expression. (H) Scoring of boundary expression of *rfng* in different conditions according to severity levels: wild type = normal expression of *rfng* in all boundaries; severity 1 = general reduction of *rfng* expression levels; severity 2 = partial absence of *rfng* expression leading to discontinuous boundaries; severity 3 = total absence of *rfng* boundary expression. Control (n=15); MO-*taz* (n=20); CRISPR-*taz* (n=30); MO-*yap1* (n=52); CRISPR-*yap1* (n=21); CRISPR-*tead1a* (n=23); CRISPR-*tead3a* (n=12). Dorsal views, anterior to the top. Scale bar: 50 μm.

Taz and Yap have been intensively studied as components of a pathway which links mechanical tension to the regulation of cell proliferation (Elbediwy et al., 2016; Gaspar and Tapon, 2014; Halder et al., 2012; Low et al., 2014). In addition, Taz and Yap have been implicated in the maintenance of stem cells or regulation of cell differentiation in specific tissues (Chen et al., 2019; Isomursu et al., 2019; Luxenburg and Zaidel-Bar, 2019; Mo et al., 2014; Varelas, 2014). Mechanical cues or other inputs lead to the translocation of Yap/Taz protein from cytoplasm to the nucleus, where they can interact with Tead family transcription factors to regulate specific gene expression. Gene expression studies have found that two tead family members, *tead1a* and *tead3*, are widely expressed in the nervous system, with segmental regulation of the level of expression (Thisse et al., 2001). To determine whether Taz acts together with these Tead family transcription factors to regulate boundary gene expression, we carried out transient Crispr-mediated knockouts. We found that knockout of t*ead1a*, but not of *tead3*, leads to a decrease in *rfng* expression (Fig.5F-H).

### Myosin regulation downstream of EphA4 regulates Taz localisation

To determine whether EphA4 signaling and actomyosin contraction acts by regulating the subcellular localisation of Taz protein, we first carried out immunostaining studies during normal hindbrain development. We found increased nuclear localisation of Taz at hindbrain boundaries, starting at 14 hpf, and becoming more prominent at 18 hpf (Fig.6A-C, G). To determine whether Taz localisation is regulated downstream of EphA4, we carried out immunostaining in *epha4* null mutants. We found that there is a loss of nuclear Taz staining at the r2/r3, r3/r4 and r5/r6 borders, whereas Taz nuclear localisation occurs at the r1/r2, r4/5 and r6/r7 borders where boundary marker expression occurs in *epha4* mutants (Fig.6D). To test whether Taz localisation is influenced by myosin phosphorylation, we carried out *mypt1* knockdown and found that this leads to an increase in the number of cells with nuclear Taz at segment borders (Fig.6E, H, I). This finding is consistent with the observation of an increased number of cells expressing *rfng* following *mypt1* knockdown. Finally, we analysed the effect of decreasing myosin II function by treating embryos with blebbistatin and found a decrease in nuclear localisation of Taz (Fig.6F).

**Figure 6:**
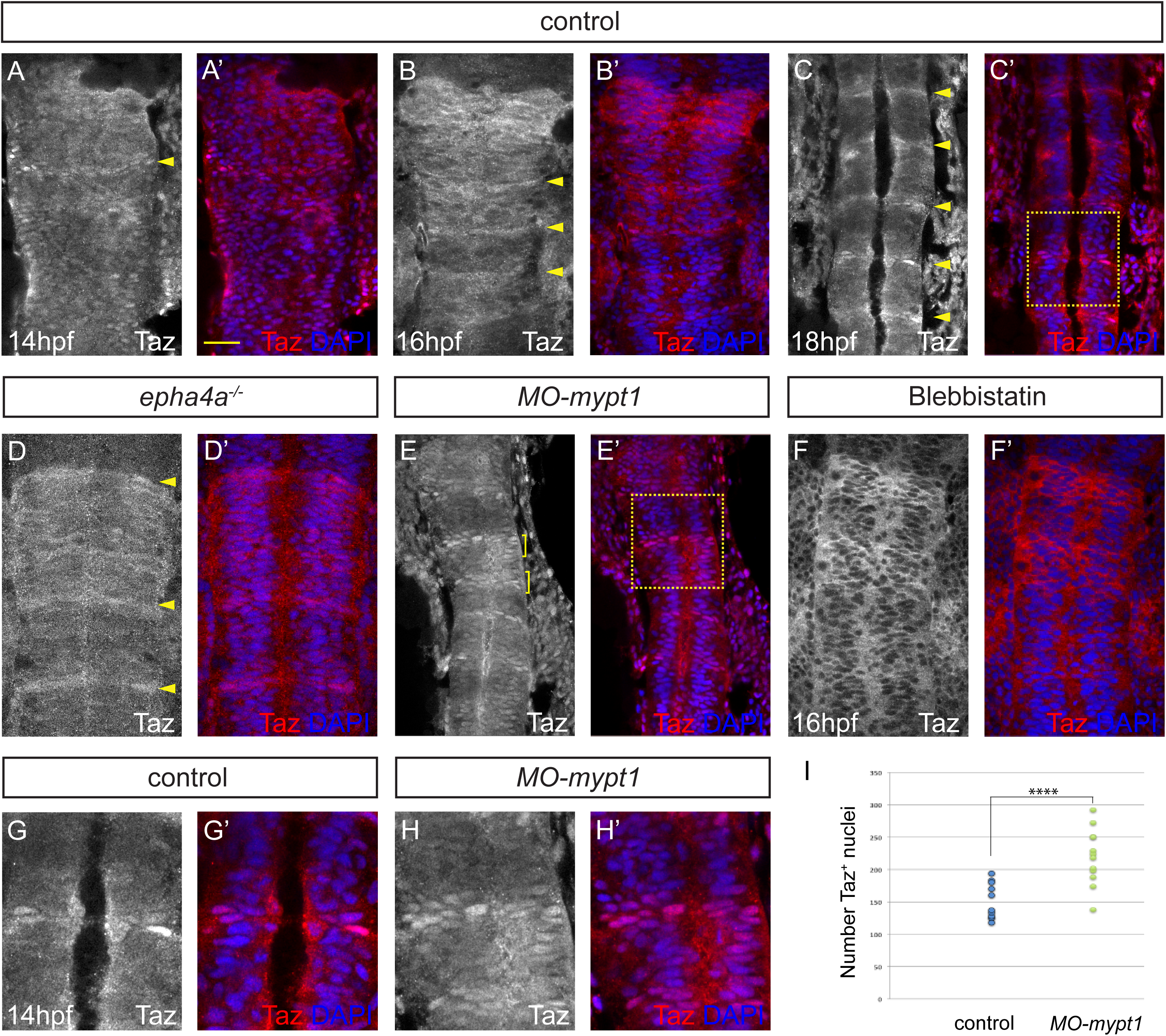
Eph-ephrin signalling and actomyosin tension regulate Taz nuclear localization. **(A-C)** Time course of the localization of Taz protein. Nuclear localization of Taz starts to be detected in hindbrain boundaries at 14 hpf (A, A’). Some boundaries have elevated nuclear Taz at 16 hpf (B, B’), and nuclear Taz is present in all boundaries at 18 hpf (C, C’). **(D**, **D’)** Nuclear Taz is reduced at specific boundaries in *epha4* mutants. **(E**, **E’)** Ectopic cells with elevated nuclear Taz are observed after *mypt1* knockdown. **(F**, **F’)** Blebbistatin treatment inhibits the nuclear accumulation of Taz at boundaries. (G, H) Higher magnification images corresponding to boxed areas in C’ and E’. (I) Quantitation of number of nuclei with Taz staining in controls (n=12) and *mypt1* knockdowns (n=13) (**** p<0.0001). Dorsal views, anterior to the top. Arrowheads indicate boundary position; brackets indicate expansion of nuclear Taz staining. Scale bar: 30 μm.

The *Drosophila* homologue of Yap/Taz, Yorkie, can increase myosin activity and tension independently of its function as a transcription co-factor (Xu et al., 2018). We therefore wondered whether Taz is required for actomyosin regulation in the hindbrain. To address this question, we analysed MLC phosphorylation following knockout of Taz, and found that pMLC is still elevated at hindbrain boundaries (Suppl. Fig.4). Taken together, these findings support a model in which EphA4 activation leads to actomyosin phosphorylation and contraction at segment borders, which in turn increases nuclear localisation of Taz and boundary marker expression.

## Discussion

A key concept that came from early studies of compartment boundaries is that sharp borders enable the correct organisation of signaling centres (Dahmann and Basler, 1999). However, it remains unclear whether or how border sharpening and boundary cell formation are coordinated. We have studied this in the vertebrate hindbrain, in which segment borders are sharpened and boundary cells form that act as a signaling centre. We show that forward signaling of EphA4, which regulates myosin light chain phosphorylation that increases cortical tension, is required both for border sharpening and for hindbrain boundary cell formation. Furthermore, increasing myosin II phosphorylation by knockdown of *mypt1* increases boundary marker expression, whereas inhibition of myosin II function or actin polymerization blocks boundary marker expression. We show that EphA4 signaling and myosin phosphorylation induce nuclear translocation of Taz, which together with Tead1a regulates boundary marker expression. Since increased tension underlies the maintenance of a straight border, cell segregation and boundary cell formation are coupled, thus ensuring that boundary cells are organised at a sharp border.

### EphA4 signaling and boundary cell formation

By generating a series of point and deletion mutants of *epha4*, we find that forward signaling is essential for boundary marker expression, with a strong input of kinase-dependent signaling and lesser input of PDZ binding domain dependent signaling. These findings are consistent with studies of the regulation of cell repulsion and cortical tension by Eph receptor signaling (Canty et al., 2017; Fagotto et al., 2013; O’Neill et al., 2016; Rohani et al., 2011; Rohani et al., 2014; Taylor et al., 2017). Cell repulsion and tension are regulated by increased Rho activity, which leads to myosin light chain phosphorylation and actomyosin contraction at borders where Eph receptor activation is occurring (Fagotto et al., 2013; Rohani et al., 2014). Multiple kinase-dependent pathways have been found to link Eph receptor forward signaling to Rho activation (Jorgensen et al., 2009; Kania and Klein, 2016; Pasquale, 2008). Eph kinase-independent signaling can also lead to cell repulsion and segregation (Taylor et al., 2017) and can activate Rho, for example through binding of Dishevelled to the PDZ domain binding motif of Eph receptors (Tanaka et al., 2003). Such kinase-independent signaling leads to a less sustained cell repulsion response than occurs when Eph kinase function is intact (Taylor et al., 2017). Taken together, these findings reveal some functional overlap between kinase- and PDZBD-dependent signaling, with a greater role of Eph kinase-activated pathways both in cell segregation and boundary cell induction.

Previous studies had not resolved whether boundary cells form on one or both sides of the interface of hindbrain segments. We find that for r3 and r5 they form on one side of each interface, in the *epha4*-expressing cells, and this is because they are induced by forward and not by reverse signaling. This finding is consistent with evidence that although reverse signaling can trigger cell repulsion, forward signaling leads to much stronger cell repulsion and actomyosin contraction and thus has a dominant role in cell segregation and border sharpening (Canty et al., 2017; Fagotto et al., 2013; O’Neill et al., 2016; Rohani et al., 2011; Rohani et al., 2014; Taylor et al., 2017; Wu et al., 2019). However, *rfng* expression is also detected in some cells adjacent to r3 or r5 that are not expressing *epha4*, in particular at the r5/r6 border. The finding that such expression adjacent to r3 and r5 does not occur in *epha4*^*Δ651*^ or *epha4*^*KD*^ mutants argues against the possibility that reverse signaling upregulates boundary marker expression. An alternative explanation is that *rfng*-expressing cells in r6 derive from intermingling of boundary cells across the segment border. This explanation requires that *epha4* expression is downregulated in r5 cells that intermingle into adjacent segments, and indeed recent work has found dynamic regulation of r3 and r5 cell identity following intermingling (Addison et al., 2018).

The decrease in boundary marker expression in *epha4* mutants is partially rescued by *mypt1* knockdown, suggesting that there is residual activation of myosin II at specific borders, perhaps due to other Eph-ephrin pairs. Intriguingly, the r5/r6 boundary was not rescued in *epha4* null mutants, but was in *epha4*^*KD*^ and *epha4*^*Δ651*^ mutants. Since the *epha4*^*Δ651*^ mutant lacks forward but not reverse signaling, whereas the *epha4* null mutant lacks both, this could suggest that reverse signaling into r6 cells can induce boundary marker expression when tension is amplified by *mypt1* knockdown. However, *rfng* expression spreads into r5 but not r6 after *mypt1* knockdown, arguing against this idea. As some EphA and EphB receptors can form heteromers (Fox and Kandpal, 2011), an alternative explanation is that truncated or kinase-dead EphA4 enables activation of another Eph receptor by ephrinB3. Indeed, EphB4, which has a low affinity for ephrinB3 (Noberini et al., 2012), is expressed in r5 and r6 and regulates cell segregation (Cooke et al., 2001).

### Regulation of cell identity by Taz activity

There is increasing evidence for roles of Yap/Taz activity in maintaining stem cells, or in some tissues in promoting their differentiation to specific derivatives (Kumar et al., 2017; Mo et al., 2014; Varelas, 2014). In some contexts, nuclear translocation of Yap/Taz protein is regulated by forces originating from interaction of cells with extracellular matrix, from stretching, shearing and compression of cells, and from actomyosin contractility within the cell (Elbediwy et al., 2016; Halder et al., 2012; Low et al., 2014; Sun and Irvine, 2016; Varelas, 2014). Hindbrain boundary cells are neural progenitors that are prevented from differentiating through Notch activation, which is promoted by Rfng (Cheng et al., 2004), thus maintaining the boundary signaling centre (Terriente et al., 2012). Activation of Taz by actomyosin contraction therefore leads to the formation and maintenance of these specialised progenitors in part through regulation of Notch pathway activity. Likewise, an interplay between Yap/Taz and the Notch pathway that maintains progenitors has been found in other tissues (reviewed by (Totaro et al., 2018)). For example, Yap/Taz maintains epidermal stem cells by inhibiting Notch signaling through regulation of Notch pathway components (Totaro et al., 2017). In another example, the contractility of muscle cells activates Yap, which upregulates Jag2 expression, leading to Notch activation in neighbours that inhibits their differentiation (Esteves de Lima et al., 2016).

Yap and Taz also have important roles in growth control in which genes that drive proliferation are upregulated by nuclear localisation of Yap/Taz, which is inhibited by activation of the Hippo pathway (Gaspar and Tapon, 2014; Halder and Johnson, 2011; Low et al., 2014). Since cortical tension leads to nuclear localisation of Taz at hindbrain boundaries, this raises the question of whether actomyosin contraction increases cell proliferation in addition to inducing boundary marker expression. Studies in chick argue against this idea as hindbrain boundaries have a lower proliferation rate than segment centres (Guthrie et al., 1991; Peretz et al., 2016), reflecting their role as a pool of neurogenic stem cells. However, recent work has found two-fold greater proliferation at boundaries than segment centres at late stages in the zebrafish hindbrain (after 26 hpf), which depends upon actomyosin and Yap/Taz/Tead activity (Voltes et al., 2018). Since this study only analysed late stages, it did not detect the role in boundary cell specification, which we find occurs prior to 18 hpf. Taken together, these findings suggest stage-specific functions of Yap/Taz activity in cell specification and proliferation at hindbrain boundaries.

### Concluding perspectives

The mechanical regulation of gene expression enables an interplay between morphogenesis and cell identity that contributes to tissue patterning (Chan et al., 2017; Kim et al., 2018; Xia et al., 2019). The transcriptional control of cell differentiation leads to differential expression of mediators of morphogenesis, creating mechanical forces which can in turn feed back on the specification of cell identity. In the hindbrain, *epha4* expression is regulated by *krox20* (Theil et al., 1998), such that cell segregation and border sharpening is coupled to segmental identity (Tumpel et al., 2009). Mechanical forces regulated by EphA4 signaling also lead to the specification of boundary cell fate, thus ensuring correct organisation of signaling centres. There is increasing evidence for roles of Eph receptors and ephrins in the regulation of cell differentiation through a diversity of pathways (Laussu et al., 2014; Wilkinson, 2014). In some cases, Eph receptor activation seems to be deployed to only regulate cell differentiation, by acting through pathways distinct from those that underlie cell segregation. For example, Eph activation regulates cell fate choices in Ciona by antagonising Fgf signaling through inhibition of the MAPK pathway (Picco et al., 2007; Stolfi et al., 2011). It will be important to understand how Eph signaling has these distinct functions in cell segregation and regulation of cell differentiation in different contexts. Since Eph signaling drives cell segregation through actomyosin regulation, and acts in many tissues, it will be interesting to determine whether it has broader roles in activating the Yap/Taz pathway to couple border formation and the control of cell identity.

## Materials and methods

### Maintenance of zebrafish strains

Zebrafish embryos were raised at 28.5°C as described (Westerfield, 2007). Embryos were staged according to morphological criteria (Kimmel et al., 1995). The zebrafish work was carried out under a UK Home Office Licence under the Animals (Scientific Procedures) Act 1986 and underwent full ethical review.

### Morpholino knockdown

Antisense morpholino oligonucleotides (Gene Tools) were injected into one-cell-stage embryos. All injections were done in *p53* homozygote mutants or in combination with a *p53* morpholino to inhibit the off-target effects mediated by activation of pro-apoptotic pathways (Gerety and Wilkinson, 2011; Robu et al., 2007). The antisense morpholinos used were a splice-blocking morpholino against *mypt1* (Gutzman and Sive, 2010) and *yap1* (Skouloudaki et al., 2009), and a translation-blocking morpholino against *taz* (Hong et al., 2005); the sequences are in Table 1. 4 ng of morpholino were injected in all cases except for MO-taz, for which 2.5 ng were injected.

**Table 1.**
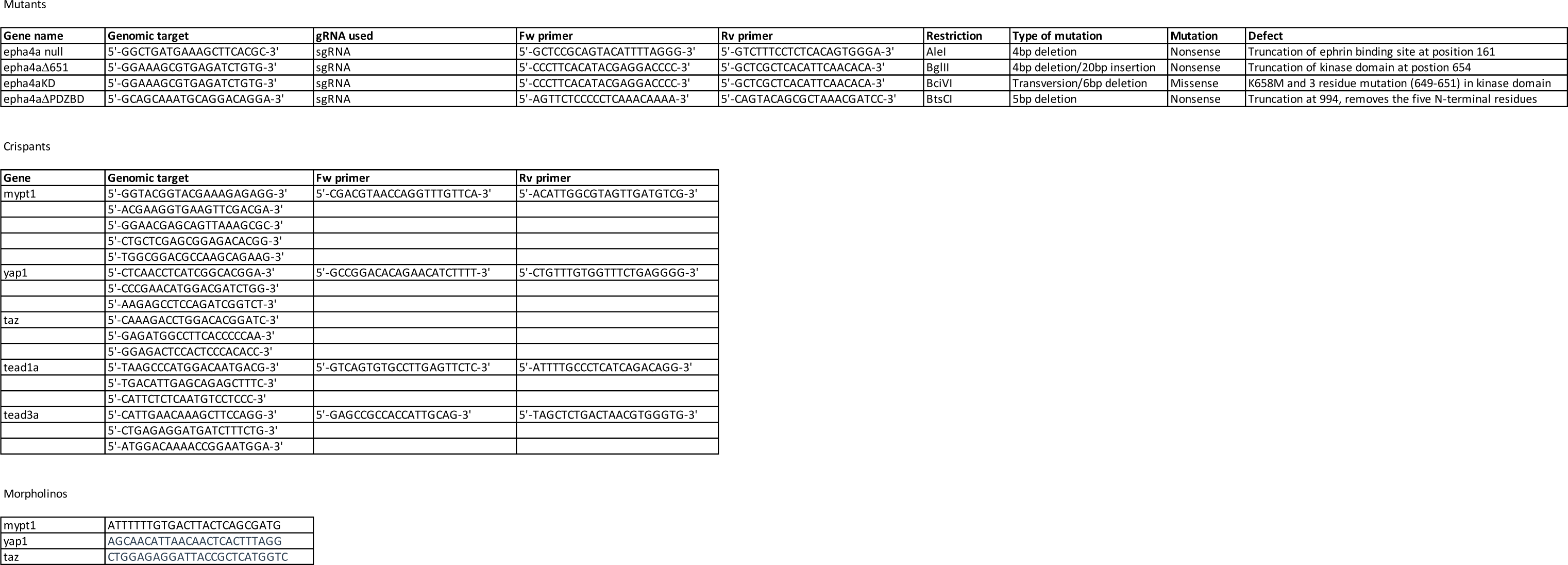

### Pharmacological treatments

Embryos were dechorionated and treated at the specified stages with 12.5 μM blebbistatin or 50 nM LatrunculinB in Danieau’s solution. Embryos were fixed in 4% paraformaldehyde (PFA) and processed for immunostaining or *in situ* hybridization.

### Generation of mutants

All injections were done in one-cell stage embryos. *ephrinb3b* mutants were generated using TALENs designed and constructed as previously outlined (Cermak et al., 2011). Plasmids used in the construction process (Golden Gate TALEN and TAL Effector Kit 1.0, #1000000016) as well as pCS2TAL3-DD and pCS2TAL3-RR destination vectors (#37275 and #37276) (Dahlem et al., 2012) were obtained from Addgene. TAL effector domains and FokI nuclease were cloned into these destination vectors to form the final pCS2-TAL vector for each TALEN, from which mRNA was synthesised using the SP6 mMessage mMachine® kit (Life Technologies). Embryos were injected with equal amounts (100 – 300 pg) of RNA encoding each of the left and right TALEN arms. A founder with a frame shift (5 bp deletion and 3 bp insertion) that truncates the protein at residue 5 was used to raise the *ephrinb3b* mutant line. RVD Sequences of *ephrinb3b* TALENs:

Left: NG HD NN NN NN NN NI NG NG NG HD NI NI NI NG NN NN HD

Right: HD NI NN NN NI NN NI NI NG NG HD HD HD NI NI NG HD HD NI NG

Point and truncated mutants of *epha4a* were generated by CRISPR/Cas9. For this, oligonucleotides targeting different *epha4a* sequences were cloned into the pDR274 plasmid for sgRNA production ((Hwang et al., 2013); #42250 Addgene). In vitro synthesis of the sgRNA was done using the T7 RiboMAX^TM^ Large Scale RNA Production System (#P1300 Promega). Embryos were injected with 200-300 pg gRNAs and 1.6 ng EnGen Cas9 protein (#M0646M NEB). The target and gRNA sequences, and mutations generated, are given in Table 1. Immunostaining for EphA4 confirmed a complete absence of protein in homozygous null embryos

To introduce the K658M mutation in the kinase domain of EphA4a, sgRNA and Cas9 protein were co-injected with a 74 bp donor oligonucleotide (AAGATGCCTGGAAA GCGTGAaATtTGcGTGGCCATAAAAACCCTAAtGGCAGGgTACACCGACAAGCAAAGGCG) containing three silent mutations at the gRNA target site, the K658M mutation and an additional silent mutation that generated an RsaI restriction site. Mutations were identified by amplicon restriction using restriction enzymes or T7 endonuclease I (#M0302L NEB) and verified by sequencing. A fish was identified carrying the K658M mutation together with a 6 bp deletion affecting 3 additional residues (649-651) in the kinase domain.

For the transient CRISPR knockouts of *mypt1, yap1, taz, tead1a* and *tead3a*, 3 to 5 crRNAs targeting the same gene (Table 1) were obtained from Integrated DNA Technologies Inc. (IDT, Iowa, USA). crRNAs were annealed with equimolar amounts of tracRNA and 100 to 150 pg of each gRNA were co-injected with Cas9 protein. The generation of deletions was validated by PCR.

### Immunohistochemistry and *in situ* hybridization

For immunohistochemistry, embryos were fixed in 4% PFA for 2 hours and processed using standard methods. For anti-Taz stainings, fixed embryos were heated at 90°C in 150mM Tris-HCl pH 9.5, rinsed and treated with DNAse1 0.025U/ml for 75 minutes at 37°C prior to staining. Samples were imaged using a Leica SP5 confocal microscope. Antibodies against Taz and pMLC were from Cell Signalling Technology (#D24E4 and 3671, respectively). Anti-EphA4 was described previously (Irving et al., 1996). Nuclear Taz staining was measured using Volocity software (Improvision) and statistical analysis carried out using unpaired two-tailed Student’s t-test.

For *in situ* hybridization, embryos were fixed in 4% PFA overnight at 4°C and kept in methanol at −20°C prior to processing. The probes used have been previously described: *egr2b* (Oxtoby and Jowett, 1993), *rfng* (Cheng et al., 2004), *wnt1* (Molven et al., 1991), *sema3gb* (Terriente et al., 2012). Digoxigenin-UTP labelled riboprobes were synthesised and *in situ* hybridization performed as previously described (Xu et al., 1994). After BCIP/NBT color development, embryos were re-fixed, cleared in 70% glycerol/PBS, and mounted for imaging using a Zeiss Axioplan2 with Axiocam HRc camera. In some experiments, *rfng, wnt1* and *epha4a* transcripts were detected by hybridization chain reaction (HCR) using reagents obtained from Molecular Instruments and the method described by (Choi et al., 2016).

## Acknowledgements

We thank Nic Tapon for discussions, and the staff of the Crick Aquatics and Light Microscopy facilities for their excellent support. This work was supported by the Francis Crick Institute which receives its core funding from Cancer Research UK (FC001217), the UK Medical Research Council (FC001217), and the Wellcome Trust (FC001217).

## Competing interests

The authors declare that they have no competing interests.

## Figure legends

**Supplementary Figure 1. Boundary marker expression in *epha4* mutants**. **(A-C)** Expression of *wnt1* is reduced at specific hindbrain boundaries in *epha4*^*-/-*^ (B) and *epha4*^*KD*^ (C) mutants compared to controls (A). Dorsal expression of *wnt1* is not changed in the mutants. **(D-F)** *sema3gb* expression is reduced at specific boundaries in *epha4*^*-/-*^ (E) and *epha4*^*KD*^ (F) mutants compared to controls (D). Arrowheads indicate normal boundary expression of *wnt1* or *sema3gb*, while stars indicate reduction or absence of boundary marker expression. Dorsal views, anterior to the top. **(G**, **G’)** HCR staining for *rfng* and *epha4a* in *epha4*^*KD*^ mutants reveals that remaining *rfng* expression occurs in *epha4*-expressing cells at the r5/r6 boundary (white arrowhead). Dorsal view, anterior to the left. Scale bar: 50 μm.

**Supplementary Figure 2. Boundary marker expression after *mypt1* knockdown**. **(A**, **B)** Expression of *wnt1* in control (A) and *mypt1* knockdown embryos (B). **(C**, **D)** Expression of *sema3gb* in control (C) and *mypt1* knockdown embryos (D). Both markers have increased expression at boundaries following *mypt1* knockdown. Dorsal views, anterior to the top. **(E**, **F)** HCR stainings for *rfng* in control (E) and a representative latrunculinB-treated embryo (F). Dorsal views, anterior to the left. Scale bar: 50 μm.

**Supplementary Figure 3. Boundary marker expression after *taz* knockdown**. **(A**, **B)** Expression of *wnt1* in control (A) and *taz* knockdown embryos (B). (**C**, **D)** Expression of *sema3gb* in control (C) and *taz* knockdown embryos (D). Boundary expression of both markers is lost following *taz* knockdown. Dorsal views, anterior to the top. Scale bar: 50 μm.

**Supplementary Figure 4. pMLC after *taz* knockdown**. **(A**, **B)** *taz* knockdown embryos immunostained to detect pMLC, shown in two different confocal planes. Increased pMLC is detected at hindbrain boundaries, suggesting that myosin phosphorylation does not require Taz function. Lateral views, anterior to the top. Scale bar: 50 μm.

