## Supplementary figures and images for "Actomyosin regulation by Eph receptor signaling couples boundary cell formation to border sharpness"

### Supplemental Figures

Supplementary Figure 1

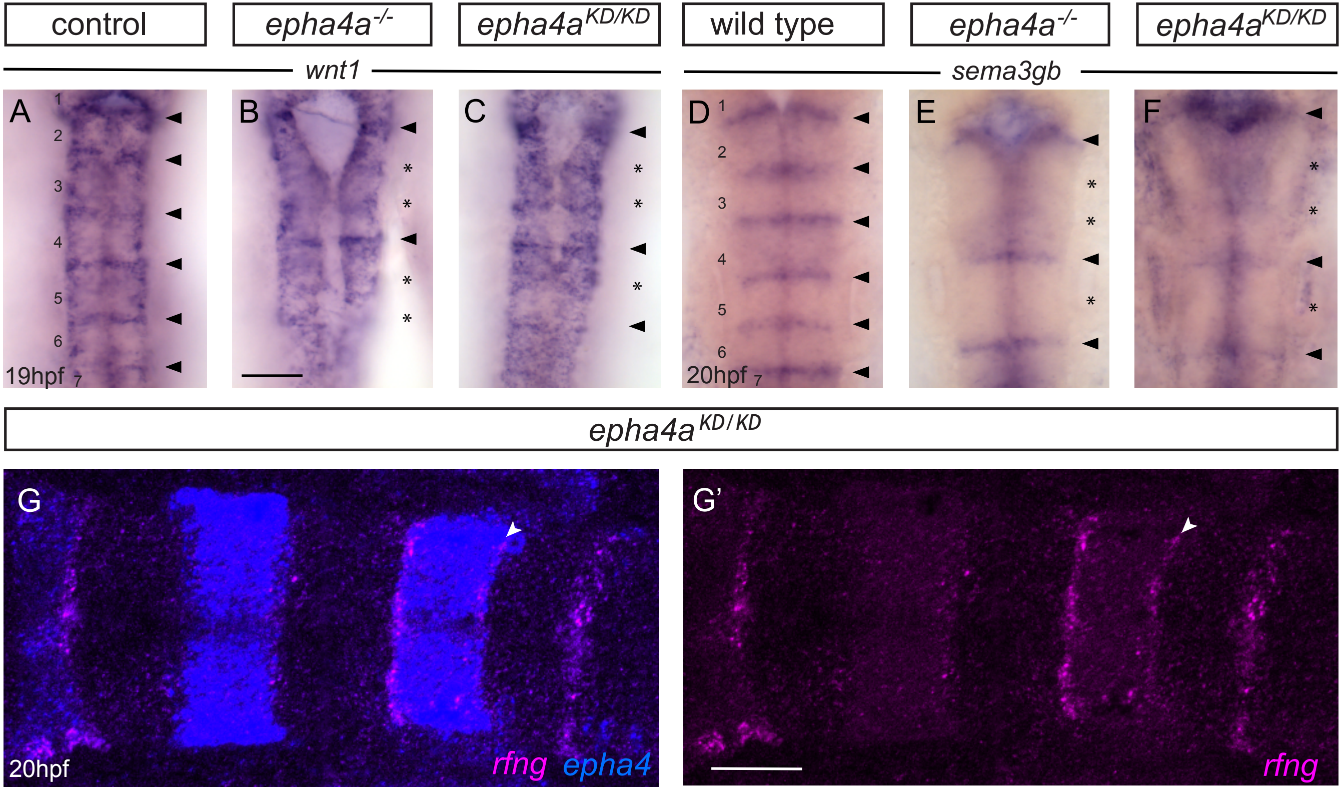

Supplementary Figure 2

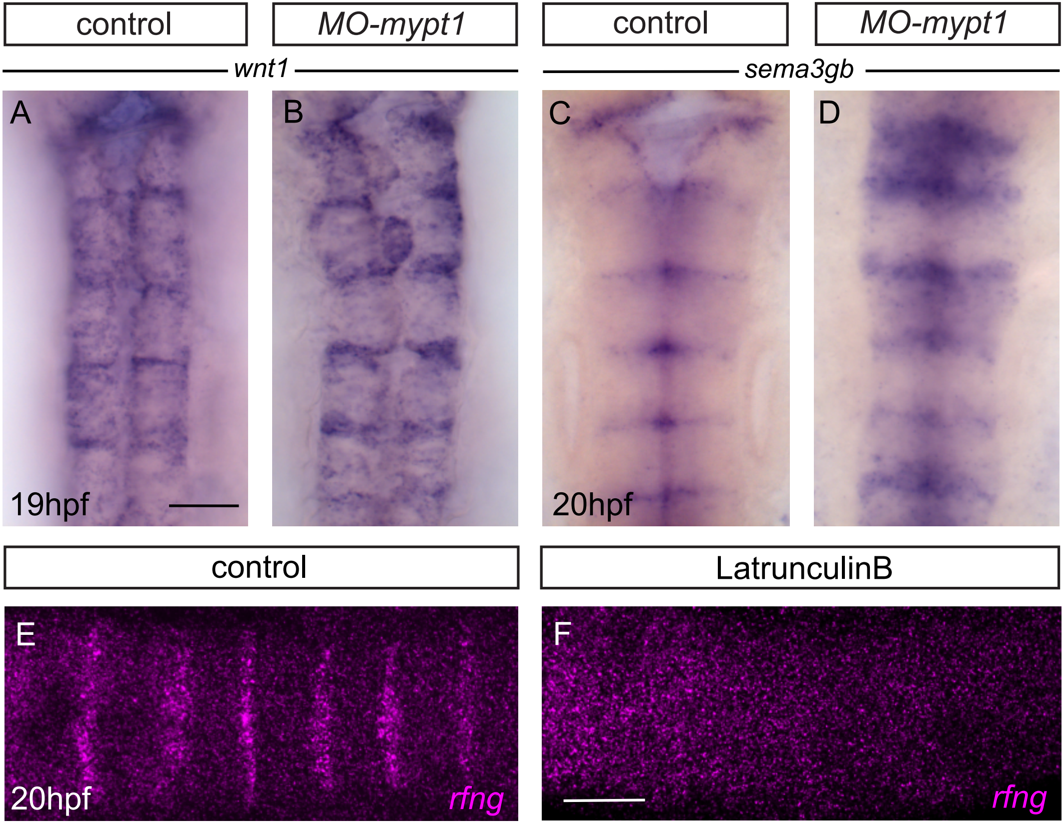

Supplementary Figure 3

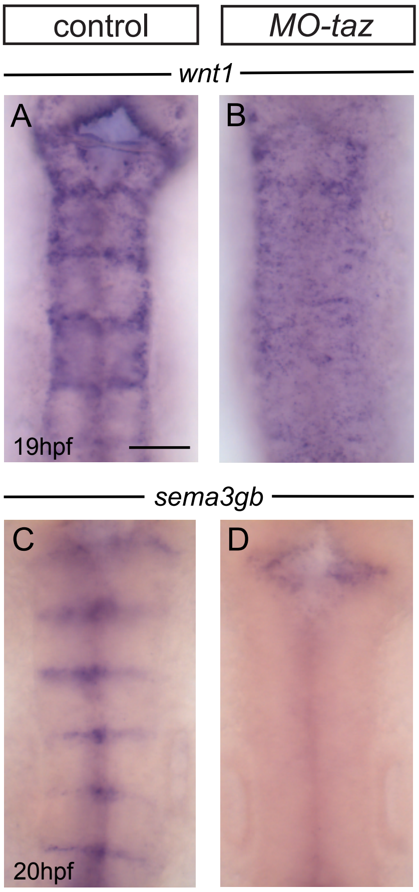

Supplementary Figure 4

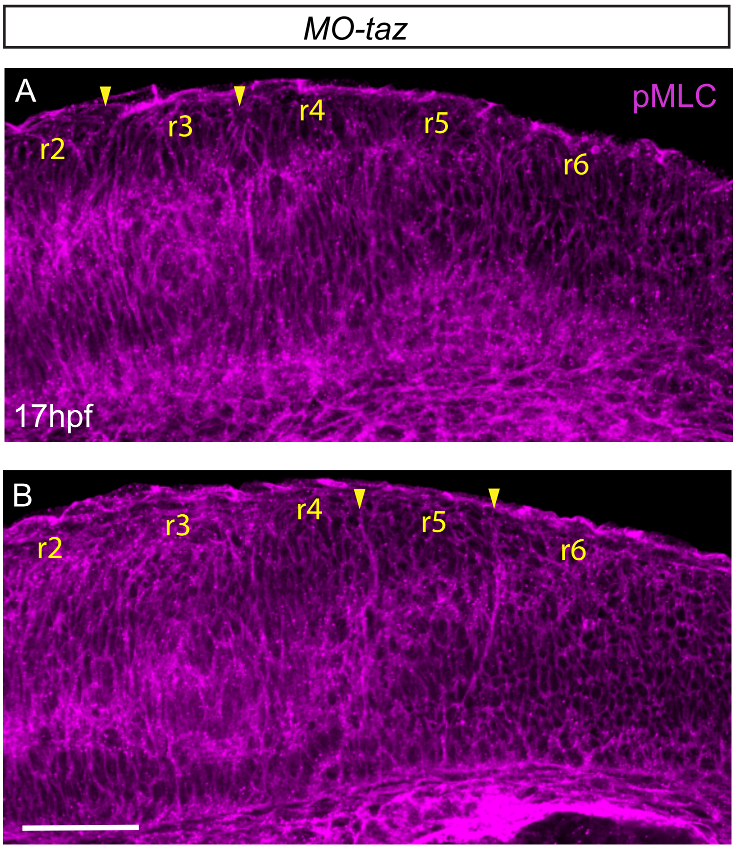
